# Joint infrared pupil images and near-corneal-plane spectral light exposure data under natural conditions across the adult lifespan

**DOI:** 10.1101/2025.03.02.641032

**Authors:** Rafael Lazar, Manuel Spitschan

**Affiliations:** Centre for Chronobiology, Psychiatric Hospital of the University of Basel, Switzerland; Research Cluster Molecular and Cognitive Neurosciences, University of Basel, Switzerland; Department of Biomedicine, University of Basel, Switzerland; Max Planck Institute for Biological Cybernetics, Translational Sensory & Circadian Neuroscience, Tübingen, Germany; TUM School of Medicine & Health, Chronobiology & Health, Technical University of Munich, Munich, Germany; TUM Institute for Advanced Study (TUM-IAS), Technical University of Munich, Garching, Germany

## Abstract

Factors influencing pupil size can be experimentally controlled or held constant in laboratory experiments, but how the pupil reflects changes in retinal input from the visual environment in real-world viewing conditions has yet to be captured in a large, age-diverse sample. In this dataset, we address this research gap by collecting data in a hybrid field-laboratory experiment (*N*=83, 43 female, age range 18–87 years) using a custom-built, wearable, video-based eye tracker with a spectroradiometer measuring spectral irradiance in the approximate corneal plane, resulting in a total of valid 29,664 recorded spectral irradiance and eye image pairs along with 83 approximately 3-minute-long calibration videos. After an initial 3-minute calibration procedure, 10-minute dark-adaptation period and a 14-minute controlled laboratory light condition, participants moved in and between indoor and outdoor environments of varying spectral irradiance for ∼25–35 minutes and performed a range of everyday tasks. This dataset may provide a basis for developing algorithms for pupil detection, processing, and prediction under natural conditions.

## Background & Summary

In humans, light exposure triggers a range of physiological processes and pathways. The pattern of environmental light is imaged through the lens on the back of the eye, the retina. The retina contains several photoreceptor classes that convert light into neural impulses. The cones and rods underlie visual function and perception – the detection and discrimination of objects, colour and motion – across dim (rods) and daylight light intensities (cones) ^1^. Cone and rod signals are integrated in the retinal ganglion cells (RGCs) ^2^. Around 25 years ago, it was discovered that some of the RGCs are intrinsically photosensitive themselves without any input from the cones and rods due to the short-wavelength-sensitive photopigment melanopsin ^3–5^. These intrinsically photosensitive retinal ganglion cells (ipRGCs) mediate several “non-visual” functions in a dose-dependent manner, including the synchronisation of circadian rhythms ^4,6^, the suppression of the endogenously produced hormone melatonin ^7,8^ and the modulation of alertness ^9^.

How much light reaches the retina depends not only on environmental illumination but also on the size of the pupil, the effective aperture of the eye’s optical apparatus ^10^. In adult humans, the size of the pupil varies from approx. 1.5 mm to 9 mm ^11^, corresponding to a difference in pupil area of approx. 1.8 mm^2^ and 64.6 mm^2^ (∼1.5 log_10_ units). Pupil size, in turn, is controlled by environmental illumination, with smaller pupils associated with brighter illumination ^12^. Pupil size is controlled by the activity of the retinal photoreceptors, with the ipRGCs playing the most important role under steady-state illumination ^13^. The cones and rods also contribute to pupil size, encoding fast changes in illumination and variation of light levels in the scotopic range ^14,15^. The short-wavelength-sensitive cones (S cones) play a particular role in pupil control as their activity dilates the pupil ^16^, consistent with an S-cone-opponent input into RGCs ^17^.

Due to the primary role of ipRGCs in controlling the pupil, conventional photometric quantities (photopic illuminance [lux] or luminance [cd/m^2^]), which reflect a joint signal from the L and M cones, are not useful. In 2018, the International Commission on Illumination (Commission de l’Eclairage, CIE) defined a series of standard spectral sensitivity functions for the retinal photoreceptors (termed “α-opic” or “alpha-opic”) ^18,19^, including the ipRGCs, which peak in the living human eye near 490 nm ^20^. These functions are being established in a variety of applications, including practical lighting design ^21,22^. Evaluating a light with respect to these novel CIE quantities typically requires the measurement of light using a spectroradiometer, yielding a spectrum from which to calculate α-opic quantities using software (e.g. CIE toolbox ^23,24^, luox ^25^, luxpy ^26^, Spectran ^27^).

Most research on the determinants of pupil size has emerged from laboratory conditions under very well-controlled parametric variations of light ^15,28,29^ including a recently published, large dataset of pupillary images in patients with glaucoma, diabetes, alcohol consumption ^30^. Under naturalistic, real-world conditions, light exposure is more variable and ill-defined than in the laboratory, a term referred to as the “spectral diet” ^31^. Generating useful evidence *for* the real world, therefore, requires measurements *in* the real world through ambulatory setups ^32–34^. In the domain of pupil measurements, such data is very scarce ^35,36^ with no open data sets with conjoint light measurements currently available. ^34^

Beyond the “low-level” influences of light on pupil size and as a convenient read-out of photoreceptor function, the pupil has also received attention for higher-level processes, including cognition and learning (see recent reviews ^37,38^). In these studies, pupil size is used as a non-invasive marker for stimulus processing and cognition. With the recent growth of literature in this field, automated, video-based methods for pupil measurements continue to be developed and refined ^39–44^. Novel algorithms can often benefit from the availability of relevant datasets containing images of pupils under a wide variety of conditions for many different observers.

Here, we present a novel data set of human pupil data for a diverse sample of observers (*N*=83, 43 females; aged 18 to 87 years) under a range of different illumination conditions as they participated in a 1-hour study protocol while wearing a head-mounted video-based infrared eye tracker (Pupil Core, Pupil Labs GmbH, Berlin, Germany) ^45,46^ along with a spectroradiometer mounted in the near-corneal plane on the forehead. The data set comprises joint measurements of pupil size (images) and spectral irradiance. The results of specific hypothesis tests based on the processed data, pupil size estimations and α-opic irradiances have been published as a Registered Report ^47^.

This data set is of interest for various reasons. First and foremost, the data presented here can be used to develop and refine novel statistical and computational models that link input (light) to output (pupil size). Such models have particular relevance for assessing light exposure and retinal illumination in occupational hazards and other practical fields. Second, the availability of high-quality images of the human pupil from a diverse pool of observers across the lifespan can facilitate the development of novel image processing and computer vision approaches for detecting pupil size under real-world conditions. Third, the spectroradiometric data reported here can be used to provide insights into the statistical properties of the “spectral diet”.

## Methods

The methods associated with this dataset, together with additional processing steps, analyses and specific hypothesis tests, are described in our previously published Registered Report ^47^. Here, we provide an extended version of the data collection methods, focusing on the raw data that comprise the current dataset ^48^.

### Recorded data

#### Spectroradiometric measurements

We recorded the irradiance spectrum near the corneal plane using a commercially available, small-sized, research-grade spectroradiometer (STS-VIS, Ocean Insight Inc., Oxford, UK; Fig. **1A**), which measures in the 350–800 nm range with a wavelength accuracy of ±0.13 nm and an optical resolution of 6 nm (100 µm slit). The instrument was factory calibrated and listed with a signal-to-noise ratio of >1500:1 and a dynamic range of 4600:1. The integration times were preset to a range of 10 µs–5 s. The STS spectrometer was equipped with a direct-attach cosine corrector (CC-3-DA, diameter: 7140 µm) with a 180° field of view.

**Figure 1.**
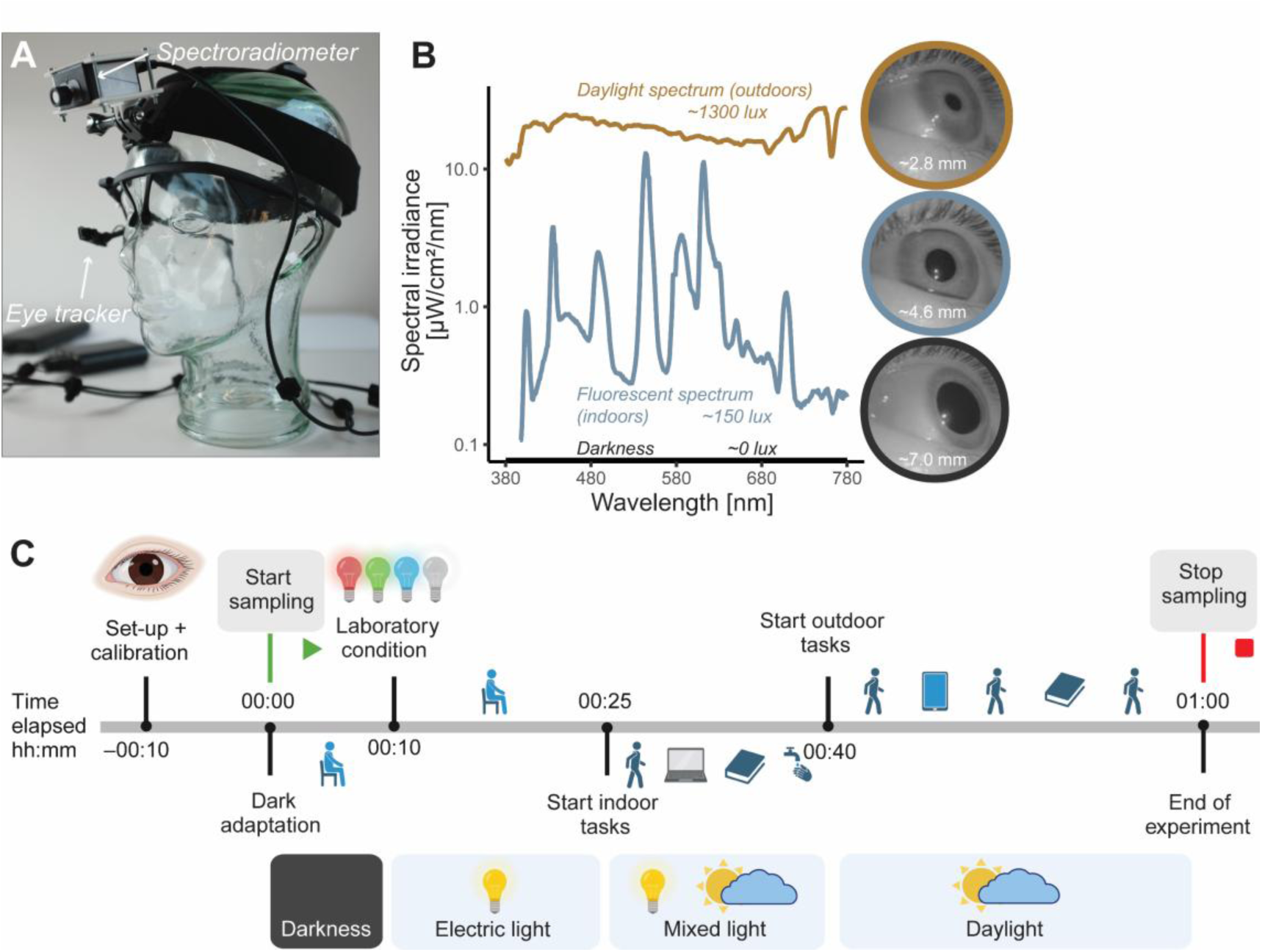
Study equipment, measurements, and protocol. **A**, The ambulatory measurement setup on a glass mannequin head. The fully portable setup consisted of an infrared-based eye tracker, a small spectroradiometer and a control computer (Raspberry Pi) together with a power bank. **B**, Examples of recorded data comprising spectral irradiance samples and corresponding illuminance values from three different lighting situations, together with pupil images and the extracted pupil sizes. The individual depicted is author R.L. **C,** Data collection protocol. After the initial eye recording for later calibration, the laboratory phase of the experiment began with a 10-minute dark-adaptation period, followed by the ∼14-minute laboratory light intervention with four spectrally different light phases at three intensities. This was followed by the field condition, which systematically included different everyday tasks and situations (walking, reading on paper and tablet, looking around outdoors, working on a laptop, washing hands, talking, etc.).

The spectral irradiance data collected by the spectroradiometer were spaced at irregular wavelengths in the visible range and thus resampled to 1 nm intervals between 380 and 780 nm, applying energy preserving interpolation. Furthermore, defect or missing pixels were linearly interpolated. The data was then corrected using the dark and thermal noise calibration, based on a polynomial model and integration time and board temperature values, measured by internal sensors (see ***Dark and thermal noise spectroradiometer calibration*** in section ***Technical Validation***). The present dataset ^48^ includes the calibrated spectral data as well as the uncalibrated CSV files (in each participant’s subdirectory “*spectra*”). The latter include the integration time and board temperature measurements, which were used in the dark and thermal noise calibration (See more details in *Data records*).

In the experiment, a customised 3D-printed case enclosed the spectroradiometer (see Fig. **1A**), which was designed and fabricated in collaboration with digitalwerkstatt GmbH in Basel, Switzerland. The case was adjustable in angle and mounted on a wearable, padded headband, which can be adapted to different head sizes. We designed the case’s connective joint to match typical action camera accessories to allow for the versatile use of low-cost replacement accessories. The template for 3D-printing the case is included in the Supporting materials uploaded to figshare ^49^. The casing was centrally located on the forehead, approximately aligned with the corneal plane and facing the direction of the observer’s gaze. The spectroradiometer lens was angled 15° below horizontal by default for all participants at the typical outdoor gaze angle ^50^. Irradiance spectra were measured every 10 seconds and saved on a portable Raspberry Pi-based microcontroller. Example spectral irradiance spectra from the recorded data, together with illuminance values, are shown in Fig. **1B**.

#### Pupil imaging

Before each experimental trial, a video-based infrared eye tracker (Pupil Core, Pupil Labs GmbH, Berlin, Germany; Fig. **1A**) was used to record an approximately 3-minute-long video of the right eye in a calibration procedure (see videos in the “*gazecalibration/000*” subdirectories of each participant’s directory, ^48^). During the calibration procedure, participants were prompted to move their eyes according to the instructions on the laptop screen ^49^. The investigator then verified that a 3D model supplied with the software by the eye tracker manufacturer (Pupil Capture v.1.15.71, Pupil Labs GmbH, Berlin, Germany) could reliably detect pupil size during different head and eye movements and positions. The 3D model ^45^ assumes that the 3D pupil can be modelled as a disc that is tangent to a rotating sphere at any instant. The Pupil segmentation is then employed to extract elliptic pupil contours, and a nonlinear optimisation procedure is used to estimate the centre of the eye sphere.

The same infrared eye tracking device was subsequently used in the experimental trials to capture still images at 10-second intervals (640 x 480 pixels, 96 dpi, jpg format) of the participant’s right eye simultaneously with the irradiance spectra recorded by the spectroradiometer. The images captured by the camera were retrieved through OpenCV and saved on the identical microcontroller as the spectroradiometric measurements. Embedded programming on the microcontroller was provided by Ocean Insight Inc. engineering. These images are saved in the same folder as the spectral irradiance data (in each participant’s subdirectory “*spectra*”) of the current dataset ^48^. Example pupil images, together with extracted pupil size in mm, are shown in Figure **1B**.

#### Study Protocol

Participants completed a variety of natural tasks in naturalistic, everyday lighting conditions (∼25–35 minutes), as well as a 10-minute dark adaptation and a ∼14-minute laboratory experimental period during a 1-hour ambulatory protocol (∼300-360 samples in total), while their approximate corneal spectral irradiance and pupil size were measured. To obtain enough valid samples per condition, each natural task and light condition lasted at least 60 seconds, given the 10-second sampling interval. Figure **1C** details the experimental protocol, including tasks, location, and light conditions. The tasks included walking, talking, reading texts on paper and tablet, looking around outdoors, filling out a questionnaire and looking up the weekly weather on a laptop, washing hands, and examining different objects. Under the uncontrolled lighting conditions, illuminance ranged from a few to several thousand lux (depending on weather conditions). The first half of the field condition was spent indoors, where participants were exposed to various light conditions from different sources, including fluorescent and incandescent room lighting, LED lighting (room lighting and laptop screens), and (indirect) daylight through the windows. The protocol then moved outdoors on a fixed outdoor route on the psychiatry campus, again involving a range of everyday situations that included mostly direct and reflected daylight and some electric light from buildings on campus. Finally, the participant was guided back to the room where the field condition started, and the measurements were stopped. All trials were conducted during natural daylight hours.

In the laboratory-based phase of the experiment (see Fig. **1C**, 00:10), the light source employed was a vertical LED front panel measuring 220 cm in width and 140 cm in height. The said panel was mounted 80 cm above floor level, and it was composed of 24 LED modules (RGB and white primaries), each with 144 LEDs. The total number of LEDs contained by the light source was 3,456, with the LED primaries’ peak emission wavelengths approximating: blue — 467 nm, green — 527 nm, and red — 630 nm under maximum intensity. The panel surface was coated with a diffuser film. Scenarios for illumination were set using DMXControl (version 2.12.2) software (DMXControl Projects e.V., Berlin, Germany). Participants were seated on a chair positioned in front of the light panel for 14 minutes (∼75 cm distance from the eyes, height varied with participant’s height, ∼90–140 cm) and were instructed to look at a cross in the centre of the panel. The sequential order of the four LED light phases (R-G-B-W) was randomised, with each phase accompanied by a 20-second dark interphase presenting a light stimulus of <0.2 lux illumiance prior to and following each light phase. Recorded samples during these dark interphases were considered “mixed” transition samples rather than “lab” phase samples. During each 3-minute phase of illumination, there were three levels of illuminance in ascending order (10 lux, 100 lux, and 250–1000 lux), each of which lasted for one minute. The “red” and “blue” conditions achieved their highest levels of illuminance at 480 lux and 250 lux, respectively, while both “green” and “white” conditions reached 1000 lux. Overall, the illumination protocol lasted 13 minutes and 40 seconds. Counterbalancing the order of the conditions resulted in 24 different sequences of the respective red (R), green (G), blue (B) and white (W) primaries, which are recorded in the laboratory log file and demographic data. Please refer to Fig. **2** for the setup during the laboratory phase.

**Figure 2.** Laboratory set-up depicted during the red LED light condition (“R”) of the laboratory phase of the experiment. The individual depicted is author R.L. The sequential order of the four LED light phases of the experimental phase (“R-G-B-W”) was randomised, with each phase accompanied by a 20-second dark interphase presenting a stimulus of <0.2 lux illuminance prior to and following each light phase. Participants were seated on a chair positioned in front of the light panel for 14 min (75 cm distance from the eyes, height varied with participant’s height, ∼90–140 cm) and were instructed to look at a cross in the centre of the panel. During each 3-minute phase of illumination, there were three levels of illuminance in ascending order (10 lux, 100 lux, and 250–1000 lux), each of which lasted for one minute. The “red” and “blue” conditions achieved their highest levels of illuminance at 480 lux and 250 lux, respectively, while both “green” and “white” conditions reached 1000 lux. Overall, the illumination protocol lasted 13 minutes and 40 seconds.

#### Descriptive data

Demographic and ancillary data were collected to describe the sample accurately. Covariates recorded comprise sex, handedness, body-mass index (BMI), vision correction, date and time of the experiment and health screening, time since awakening, sleep duration, habitual and acute caffeine consumption, chronotype, acute sleepiness, iris colour and weather conditions. Except for the final three, these covariates were obtained through a survey completed before and during the protocol.

- The experimenter rated the colour of the iris and the weather conditions. Iris colour was evaluated using a 3-point Likert-scale item, categorised as “brown”, “hazel-green”, and “blue” ^51^, before the protocol began. Weather conditions were rated at the commencement of the outdoor phase of the protocol (see Fig. **1 C**, 00:40) using a custom 4-point Likert-scale item, including the choices of “overcast”, “cloudy”, “partly cloudy”, and “sunny”.
- Acute daytime sleepiness was assessed using a German paper-pencil version of the Karolinska Sleepiness Scale ^52^, both during the indoor tasks (see Fig. **1C**, 00:30) and at the end of the protocol. The 9-point Likert-scale item ranges were labelled on odd steps.
- Both the habitual and acute caffeine consumption questionnaires were developed based on median values of caffeine content ranges in mg for typical caffeinated beverages and foods ^53^. Each of the two questionnaires consisted of 15 items on beverages, two on chocolate, and one on caffeine pills. Each item required the participant to specify how many portions of the caffeinated item they consumed, using a scale that ranges from zero (no portion) to eight portions. Each item portion corresponds to a standard serving size in ml or g and includes a caffeine content in mg. The “habitual” questionnaire asks about caffeine intake on an average day, while the “acute” questionnaire focuses on the last six hours. For habitual and acute intake, each participant’s caffeine intake was quantified as an absolute sum and a relative sum value in mg (divided by weight in kg). The questionnaires were completed as part of a “laptop task” during the protocol (∼00:30 in Fig. **1C**).
- Chronotype was assessed using a German version of the µMCTQ, an abbreviated version of the Munich ChronoType Questionnaire ^54^. The questionnaire comprises six fundamental questions, enabling a fast and effective assessment of chronotype. The mid-point of sleep on work-free days corrected by potential sleep debt (MSF_SC_) was computed as the indicating value for chronotype. Sleep duration and time elapsed since waking up are determined by answers to four open-ended questions. Participants report their bedtime and wake-up time for the night before the experiment, as well as any naps taken on the days leading up to and including the experimental day. Sleep duration was calculated by subtracting the wake-up time from the bedtime and adding the duration of naps. Time elapsed since waking up is determined by subtracting the wake-up time from when participants began filling out the questionnaire. Both the µMCTQ and the queries about sleep length and duration after waking up were completed at the start of the protocol as a component of a “laptop task” (∼00:30 in Fig. **1 C**).
- Sex, handedness, BMI (calculated from weight and height) and visual correction (“no correction”, “minus correction”, or “plus correction”) were assessed via simple demographic items as part of the survey completed just before the start of the protocol.

#### Recruitment of participants

A study flowchart illustrating the recruitment process is shown in Supplementary Figure 1 of the Registered Report ^47^, and details are also described in the methods of that article. Following our recruitment plan from the approved Stage 1 registration, data were collected until we reached our resource limit. The study was advertised with placards and flyers at the University of Basel as well as via the website of the Centre for Chronobiology (www.chronobiology.ch), on the web page of the UPK (https://www.upk.ch/) and on Unimarkt Universität Basel (https://markt.unibas.ch/). Full participation in the study was compensated with 30 Swiss francs (CHF). All study activities, including the health and eligibility screening and the experimental trial, were conducted in the Psychiatric Hospital of the University of Basel (UPK), Switzerland. The screening, trial preparation and indoor activities of the experimental trial were conducted at the Centre for Chronobiology of the UPK Basel, while all outdoor activities were conducted on the campus of UPK Basel. Mandatory measures to prevent the spread of COVID-19 in Switzerland were implemented during data collection. *N*=113 volunteers were invited to participate in the eligibility and health screening after giving written informed consent. Seven of the invited volunteers did not meet the inclusion criteria, five due to the visual tests and two due to health reasons and medication with a potential influence on the regulation of their pupils. The protocol was allocated to the remaining 106 subjects. *N*=87 completed the trials without equipment problems, while data from *n*=19 subjects were invalid due to technical problems (loose cables, camera slippage or software/format errors) and had to be excluded. A further four participants were then removed because more than 75% of their pupil data were of insufficient quality, leaving *N*=83 for the final dataset.

#### Participant characteristics

Only participants wearing contact lenses or requiring no vision correction were included, as our measurement setup was incompatible with spectacles. All participants had to achieve a visual acuity of at least 20/40, as assessed by the Snellen chart ^55^, and normal colour vision, as assessed by the HRR test ^56^. Furthermore, relevant medication, neurological and metabolic disorders, eye conditions, and mental health (assessed through the German version of the GHQ-12 ^57^ were screened in participants. In addition, the consumption of drugs (urine-based multi-panel drug test, nal von minden; Moers, Germany) and alcohol (saliva-based alcohol test, ultimed; Ahrensburg, Germany) were tested, and participants were only included when they tested negative. Table **1** shows the self-reported and experimenter-rated characteristics of the included participants. The final sample of *N*=83 participants (43 female, 40 males; *M_Age_*=35.70, *SD_Age_*=17.16; *M_BMI_*=22.96, *SD_BMI_*=3.47) was slightly biased towards younger ages, with five individuals over the age of 64 years and 30 individuals between the ages of 18 and 24 years. The experiment included 40 participants in the summer (48%), 20 in the winter (24%), 16 in the autumn (19%), and 7 in the spring (8%). Weather conditions varied between light rain (11%), very cloudy (19%), cloudy (17%), somewhat cloudy (13%), and sunny (40%). The different weather conditions during the trial were linked to distinct light level distributions (see Fig. 3 in the Registered Report ^47^). Table **1** provides further details on the participant characteristics, including chronotype, sleep duration, experiment start time, and caffeine intake. The demographic data of the included 83 participants are part of the current dataset (demographic_data.csv, ^48^). Detailed demographic data of all 113 invited individuals (including single items) are contained in the analysis code and processed data of the Registered Report ^47^, available on GitHub ^58^.

**Figure 3.**
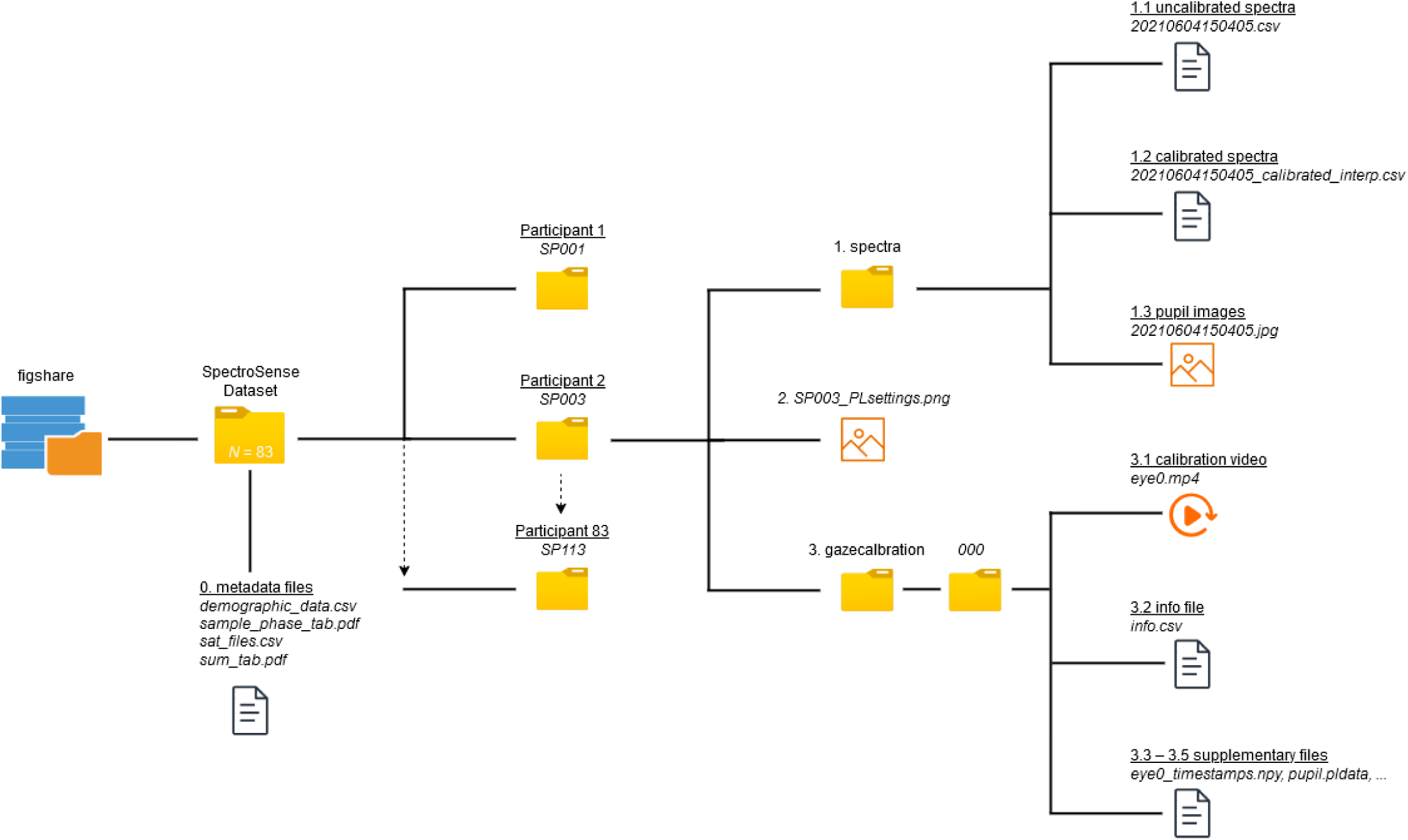
Directory structure of the data records on figshare. The *SpectroSense Dataset* is a (zip) folder with 83 subdirectories (*SP001* to *SP113*), one for each participant ^48^. The dataset is supplemented by a set of four metadata files. Each participant’s folder contains the following content: 1. The *spectra* folder, which contains the main data (uncalibrated spectral irradiance samples, calibrated spectral irradiance samples, and pupil images, 2. The *PLsettings* file (e.g. “SP003_PLsettings.png”) showing a screenshot with metadata of the pupil extraction from the images, 3. The *gazecalibration* subdirectory containing the eye calibration video, which was used for generating the 3D model for pupil size extraction.

**Table 1.** Demographic characteristics of the included participants. Age, sex, handedness, visual aid status, body mass index (BMI), caffeine consumption, sleep duration, sleep timing and sleepiness during and after the trial were self-reported. Iris colour and weather during the test were assessed by the experimenter. Season was derived from the date of testing. Numerical variables are presented as “mean (standard deviation)” and categorical variables as “count (%)”.

| Variable | Valid data, $n=83$<sup>†</sup> |
| --- | --- |
| Age [years] | 35.70 (17.16) |
| Age group [years] |  |
| 18-24 | 30 (36%) |
| 25-34 | 18 (22%) |
| 35-44 | 13 (16%) |
| 45-54 | 9 (11%) |
| 55-64 | 8 (9.6%) |
| >64 | 5 (6.0%) |
| Sex (assigned at birth) |  |
| Female | 43 (52%) |
| Male | 40 (48%) |
| Handedness (self-report) |  |
| Right | 75 (90%) |
| Left | 7 (8.4%) |
| Both | 1 (1.2%) |
| Uses visual aid |  |
| No | 57 (69%) |
| Yes - correction for myopia | 18 (22%) |
| Yes - correction for hyperopia | 8 (9.6%) |
| Wearing contact lenses during trial |  |
| No | 67 (81%) |
| Yes | 16 (19%) |
| Body Mass Index (BMI) [kg/m<sup>2</sup>] | 22.96 (3.47) |
| Midsleep on free days corrected for oversleep [h] | 2.36 (3.27) |
| Hours of sleep before trial [h] | 7.92 (2.48) |
| Average sleep duration [h] | 8.28 (2.37) |
| Estimated acute caffeine consumption [mg] | 66.55 (80.59) |
| Estimated habitual caffeine consumption [mg] | 174.76 (172.64) |
| Iris colour (experimenter-rated) |  |
| Blue | 25 (30%) |
| Hazel/green | 17 (20%) |
| Brown | 41 (49%) |
| Weather during trial (experimenter-rated) |  |
| Light rain | 9 (11%) |
| Very cloudy | 16 (19%) |
| Cloudy | 14 (17%) |
| Somewhat cloudy | 11 (13%) |
| Sunny | 33 (40%) |
| KSS-assessed sleepiness (during trial) | 3.73 (1.87) |
| KSS-assessed sleepiness (after trial) | 3.19 (1.59) |
| Season |  |
| Spring | 7 (8.4%) |
| Summer | 40 (48%) |
| Autumn | 16 (19%) |
| Winter | 20 (24%) |

<sup>†</sup> Mean (SD); $n$ (%)

#### Known limitations of the dataset

We are aware that our data is limited in the following ways:

- *Top-down determinants of pupil size*. In this work, we have not taken into account other determinants of pupil size, such as fatigue ^59^, attentional processes ^60,61^, recognition ^62^, high-level image content ^63^, target detection ^64^, mental effort ^65^, and arousal ^66^. These top-down effects were expected to be transient and relatively small compared to the light and age effects ^11,28^. In addition, we have omitted the effects of viewing distance, which affects pupil size through a common neural pathway due to the near triad of convergent eye movements and ocular accommodation (see review ^10^).
- *Estimating retinal irradiance under natural viewing conditions*. In this study, we used a spectral irradiance sensor close to the corneal plane, placed on the forehead and adjusted to a typical outdoor viewing angle (15° below horizontal, ^50^). “Shielding” effects of the eyelid under bright light conditions ^67^ and other limitations of the visual field ^68^ were not explicitly simulated with the measurement setup of this study. Given the geometry of the sensor and the human head, this was the closest we could get to feasibly measuring corneal light exposure.
- *Omission of scene radiance*. In the present dataset, we measured spectral irradiance rather than spectral radiance. This neglects the effects of variations in light levels across the field of view in different lighting scenes. Given the limitations of the real-world setting, the technical characteristics and calibration of the available instruments, it was not feasible in the present study to collect real-world radiance data and artificially limit the angular acceptance of our spectroradiometer.
- *Light history.* In the present data collection, we did not record prior light exposure (e.g., before the experiment). It is, therefore, possible that our data may contain carry-over effects from light exposure before the experiment. To provide some control during the experiment but retain the generalisability in the field recordings, we counterbalanced the light phases in the laboratory part of the experiment while keeping constant the sequence of the rest of the protocol (dark adaptation and field experiment procedure).
- *Dark and thermal noise*. Spectroradiometric measurements are affected by noise, which was taken into account by implementing a dark and temperature noise calibration described in the ***Technical Validation***. Nevertheless, spectral irradiance measurements are still affected by some baseline noise, especially when the integration time of the spectroradiometer is limited and measurements are taken in darkness. In the present study, the integration time of the spectroradiometer was limited to 5 seconds to meet the 10-second sampling intervals. Spectral irradiance samples taken in darkness may, therefore, erroneously still contain values up to ∼1.5 lx (illuminance).
- *Temporal resolution.* We likely did not capture some of the faster dynamics of pupil size regulation that may arise from cone signalling pathways ^14,16,69^, given our temporal sampling resolution (10-second intervals). Our rather low temporal resolution does not allow a detailed analysis of the fast-responding aspects of pupil size regulation or the rate of adaptation.
- *Exhaustive lighting scenarios.* We have limited our design to lighting scenarios that balance feasibility with generalisability, as the number of possible lighting scenarios is non-exhaustive. This generalisability remains conceptual, as the “true” statistical characteristics of the human spectral diet are unknown.

## Data Records

All data described here are made available using figshare ^48^ in the data structure as detailed below. Previously, corresponding processed data ^70^ were published as part of a Registered Report ^47^, together with general supporting material for the experiment conducted ^49^.

The data recording procedures described in the ***Methods*** section produced a dataset of spectral irradiance samples combined with concurrent pupil images, recorded in 10-second intervals. These data are the focus of the dataset and current data descriptor publication. The complete “*SpectroSense Dataset*” is organised in a structured folder system. Data were collected from 83 participants (first: *SP001*; last: *SP113*), with data from each participant stored in individual folders. In total, the dataset contains 29,664 recorded spectral irradiance and eye image pairs, along with 83 approximately 3-minute-long calibration videos and further contextual data. The folder structure and its contents are described below and illustrated in **Fig 3**. Please note that the valid (corrected) timestamps of each sample are contained in the filenames of the calibrated and uncalibrated spectral irradiance samples and pupil images.

## 0. Metadata files

Four metadata files are included in figshare to support using the main data given in the *SpectroSense Dataset*.

0.1 **demographic_data.csv** contains the main demographic and ancillary data recorded before and during the trial for each participant (see *Descriptive data* in *Methods*). This includes the date of participation (*date*), start time of the health screening (*begin*), age (*age*) and age group (*age_group*) in years, sex assigned at birth (*sex*), handedness (*handedness*), visual aid status (*visual_aid*), body mass index (*BMI*) in kg/m², chronotype based on mid-sleep time (*MSFsc*), time since awakening in hours (*time_awake*), sleep duration in the night before the trial in hours (*sleeping_hours*), average weekly sleep duration in hours (*SDweek*), absolute sum of acutely reported caffeine consumption in mg (*acute_sum*), sum of acutely reported caffeine consumption relative to weight in in mg/kg (*acute_sum_rel*), absolute sum of habitually reported caffeine consumption in mg (*habitual_sum*), sum of habitually reported caffeine consumption relative to weight in in mg/kg (*habitual_sum_rel*), expertimenter-rated iris colour (*iris_colour*), weather during the field experiment phase (*weather*), KSS sleepiness rating (1–9, with 9 being most sleepy) in the indoor part of the trial (*kss_pre*), KSS sleepiness rating (1–9, with 9 being most sleepy) after the outdoor part of the trial (*kss_post*) and season during the participant in the experiment (*season*).
0.2 **sat_files.csv** contains a list of all saturated/erroneous spectrometer samples (after rapid dark-to-light changes, 410 samples in total). The saturated spectrometer samples are still included in the dataset as placeholders for completeness. For the calibrated and interpolated spectra, the samples take the spectral irradiance value “0” at all wavelengths and should be considered as missing and excluded when analysing the data.
0.3 **sample_phase_tab.pdf** shows a table that summarises for every participant which filenames have been taken under which experiment phase condition (“dark”, “laboratory”, “field” with transition samples being considered “mixed”). Please note that spectral irradiance measurements in the “dark” condition (taken in complete darkness) are affected by baseline noise as integration time was limited to 5 seconds, which can result in these spectral irradiance samples having values corresponding to up to ∼1.5 lx (illuminance).
0.4 **sum_tab.pdf** shows a table that summarises for every participant, sex, age, the sequence of laboratory light conditions and the number of observations (pupil image and spectral irradiance pairs) in total and divided into experiment phase conditions (“dark”, “laboratory”, “field”, “mixed”).

### SpectroSense Dataset

The data in the ***SpectroSense Dataset*** folder are divided into participant-specific subdirectories. Each experimental session with a participant is stored in a participant-specific folder (*SP001* to *SP113*), which contains the following subdirectories and files:

1. The subdirectory ***spectra*** contain the main data of this dataset, namely the spectral irradiance and pupil image samples collected at 10-second intervals.

1.1 **Raw, uncalibrated spectroradiometer samples** (format: YYYYMMDDHHMMSS.csv, e.g. 2*0210618104421.csv*). The filenames contain the valid, corrected timestamps of each sample. These uncalibrated spectral light measurements include the following information:

1.1.1. *Time:* The time and date of the recording listed in the uncalibrated CSV files are incorrect due to an issue with the internal time of the Raspberry Pi Computer used for sampling. However, as mentioned above, the corrected date and time are given in the filenames.
1.1.2. *Serial Number*, *Integration time* (microseconds), and *Sensor temperature* (°C) give contextual information on the spectroradiometer device. The latter two were used to correct for dark and thermal noise (see section ***Dark and thermal noise spectroradiometer calibration*** under ***Technical validation***).
1.1.3. Spectral irradiance measurements with irregular wavelength spacing (between 337 and 822 nm) are provided as *Wavelengths* (nm), *Dark* (counts), and *Intensity* (counts). These data are not interpretable without Ocean Insight-specific calibration. Spectral irradiance data from this dataset should be analysed using the calibrated and interpolated spectroradiometer samples (see below).
1.2. **Calibrated and interpolated spectral irradiance samples** (format: YYYYMMDDHHMMSS_calibrated_interp.csv, e.g. 2*0210618104421_calibrated_interp.csv*): The CSV files contain two columns: “*wavelength*” in nm given in the visible range (380–780 nm, 1 nm spacing) and “*measurement*” containing the corresponding irradiance given in W/m². Note that these spectra were resampled to 1 nm intervals between 380 and 780 nm, applying energy-preserving interpolation and linear interpolation for missing or defective pixels. The filenames contain the valid, corrected timestamps of each sample.
1.3. Pupil images: Still images of the participant’s pupil (jpg format: 640 x 480 pixels, 96 dpi) taken with the “eye0” camera, corresponding to each spectroradiometer sample at 10-second intervals. The filenames contain the valid, corrected timestamps of each sample.
2. ***<Participant ID>_PLsettings.png.*** Each participant folder contains a png file (e.g. “SP001_PLsettings.png”) showing a screenshot of the pupil size extraction process in the Pupil Capture software ^71^. The screenshot provides metadata of the pupil extraction from the images, conducted to generate the processed data analysed in the Registered Report ^47^.This includes a visualisation of the 3D pupil model ^45^ including the *pupil intensity range*, which gives “*the minimum darkness of a pixel to be considered as the pupil*” ^72^. This pupil intensity range value took the value 10 at default and was adjusted (minimised) for every participant’s data so that the pupil was always fully covered while having as little leakage outside of the pupil as possible. Differences between participants occurred mainly due to differences in iris colour and differences in illumination of the eye during the field condition. The screenshot also contains “*Pupil min*” (10) and “*Pupil max*” (100) and “*Model sensitivity*” (0.9970) values, which were kept consistent across participants.
3. The subdirectory ***gazecalibration/000/*** contains the video-based eye-tracking calibration data recorded with Pupil Labs software. Files include:

3.1. **eye0.mp4**, **eye0_timestamps.npy**: Eye video files of the participant’s right eye during the calibration procedure involving different eye movements. The video was later used with offline pupil detection to calibrate the 3D model of the eye for analysing the pupil images.
3.2. **info.csv**: Participant session-specific metadata file containing recording information for each session. “*Recording Name*”(e.g. *SP001*), “*Start Date*” (format: DD.MM.YYYY), “*Start Time*” (format: HH:MM:SS), “*Start Time (System)*” (format: UNIX epoch time in sec), “*Start Time (Synced)*”(format: sec relative to the start of the Pupil Capture software’s internal time base), “*Recording UUID*”, “*Duration Time*” (format: HH:MM:SS), “*World Camera Frames*” (“0”) and *World Camera Resolution* (“1280×720” in pixels), “*Capture Software Version*” (1.15.71), “*Data Format Version*” (1.15.71), and finally “*System Info”* giving information on the Computer that the software was run on.
3.3. **gaze.pldata, gaze_timestamps.npy, pupil.pldata, pupil_timestamps.npy**: Supplementary files containing time-series data on pupil detection gaze and ensuring structural completeness and compatibility of the directory with Pupil Labs software ^71^ (recorded with version 1.15.71).
3.4. **world.intrinsics, world_timestamps.npy**: These files act as a placeholder to ensure structural completeness and compatibility of the directory with Pupil Labs software (recorded with version 1.15.71) but do not contain any meaningful data.
3.5. **user_info.csv**: This placeholder CSV file ensures structural completeness and compatibility of the directory with Pupil Labs software (recorded with version 1.15.71).

### Supplemental code for the current dataset

Python code used to correct the timestamp-based filenames, MATLAB code to generate the dark and temperature noise calibration, and R code to generate the metadata files for the raw dataset ^48^ as well as figures and tables of the present data descriptor are publicly accessible under the MIT licence in a GitHub repository ^73^ published together with this data descriptor.

### Processed data and laboratory log

The processed data used in the Registered Report ^47^, as well as a laboratory log including all 113 individuals invited to participate in the study (*N*=83 included), are provided in a previously published figshare repository ^70^. The latter repository includes the following:

1. The processed data (pupil diameters and alpha-opic irradiances corresponding to the raw data; cleaned, corrected, quality checked and categorized) from the *N*=83 included participants in CSV and RDA (R data) format. This dataset also includes detailed demographic information of these participants.
2. The laboratory log file comprising metadata for all invited participants (*SP001*– *SP113*). All files in the repository are explained in detail in the description of the repository ^70^.

### Analysis code for processed data

The analysis code used to analyse the processed data of the Registered Report ^47^ is available on GitHub ^58^. The code repository includes a detailed workflow description of the R code-based data analysis and hypothesis testing. It also includes detailed demographic data on all 113 invited individuals.

### Other supporting materials

Other supporting documents related to the protocol of the present raw dataset were published in another figshare repository ^49^. That repository includes the following:

1. The script used for the gaze calibration was performed before the start of each trial (format: OpenDocument Presentation Document; ODP).
2. Information about the laboratory-based part of the experiment.
3. Screenshots of the default settings used in the Pupil Labs software (Pupil Labs GmbH, Berlin; release version 1.15.71).
4. The 3D printed spectroradiometer case (upper and lower module) as tilted in the experiment (15° below horizontal angle).
5. The reading tasks used in the experimental protocol.
6. The research project ethics application (latest amendment) including informed consent and study information, and the approval by the responsible ethics commission.

## Technical validation

### Pupil model calibration

As described in the Methods section, the video-based infrared eye tracker (“Pupil Core”, Pupil Labs GmbH, Berlin, Germany; Fig. **1A**) used for taking the pupil images during the trial was also used to record an approximately 3-minute-long video of the right eye in a calibration procedure. In this procedure, the participants were prompted to follow the German instructions and targets presented on the Laptop screen, run with a ODP script (format: OpenDocument Presentation Document, see file in ^49^). This included moving their eyes and head in a systematic procedure so that the eye was recorded in very different positions and from different angles. The investigator then verified that a 3D model supplied with the software by the eye tracker manufacturer (Pupil Capture v.1.15.71, Pupil Labs GmbH, Berlin, Germany) could reliably detect pupil size during different head and eye movements and positions. The 3D model ^45^ assumes that the 3D pupil can be modelled as a disc that is tangent to a rotating sphere at any instant. The Pupil segmentation is then employed to extract elliptic pupil contours, and a nonlinear optimisation procedure is used to estimate the centre of the eye sphere. The recorded calibration videos are included in the “*gazecalibration/000*” subdirectories of each participant’s directory of the present dataset ^48^.

### Cross-device comparison for light measurements

To verify the near-corneal light exposure recorded by the small research-grade spectroradiometer used in the present dataset (STS-VIS; Ocean Insight), it was compared with a lightweight (∼27 g) “light dosimeter” (*lido*) device developed at the Lucerne School of Engineering and Architecture ^74^ to estimate near-corneal alpha-optic irradiances during field studies, comprising six channels in the visible spectrum (380– 780 nm). For cross-comparisons, the *lido* device was attached to the side of the measurement setup, on the left temple of the eye tracking device, to measure alpha-optic irradiances at 10-second intervals during the protocol in a small subset of participants (*n*=3, *SP025*, *SP029* and *SP033*) alongside the Ocean Insight spectroradiometer (see Fig. **4A**). Due to the limited dynamic range of the *lido* device and following the recommendations of the authors ^74^, values below 5 lx, including the 10-minute dark adaptation at the start of the study, were excluded from this analysis. The melanopic irradiances derived from the spectra recorded by the Ocean Insight spectroradiometer were matched to the melanopic irradiance estimates of the lido in the closest temporal proximity (max. 5-second difference). The data were then binned into 1-minute averages of melanopic irradiance and compared by Pearson correlation. Figure **4A** shows the arrangement of the two instruments compared in the measurement setup, while Figure 4 **B** shows the binned variations of melanopic irradiance measured during one study protocol (*SP033*) from both instruments, and Figure **4C** illustrates the correlation (*r*=0.92, *p*<0.001, Pearson coefficient) between the instruments for the binned data during the three recorded sessions, indicating that the measurements from the two instruments co-varied strongly, with comparable results, despite slight differences in the position and timing of the samples.

**Figure 4.**
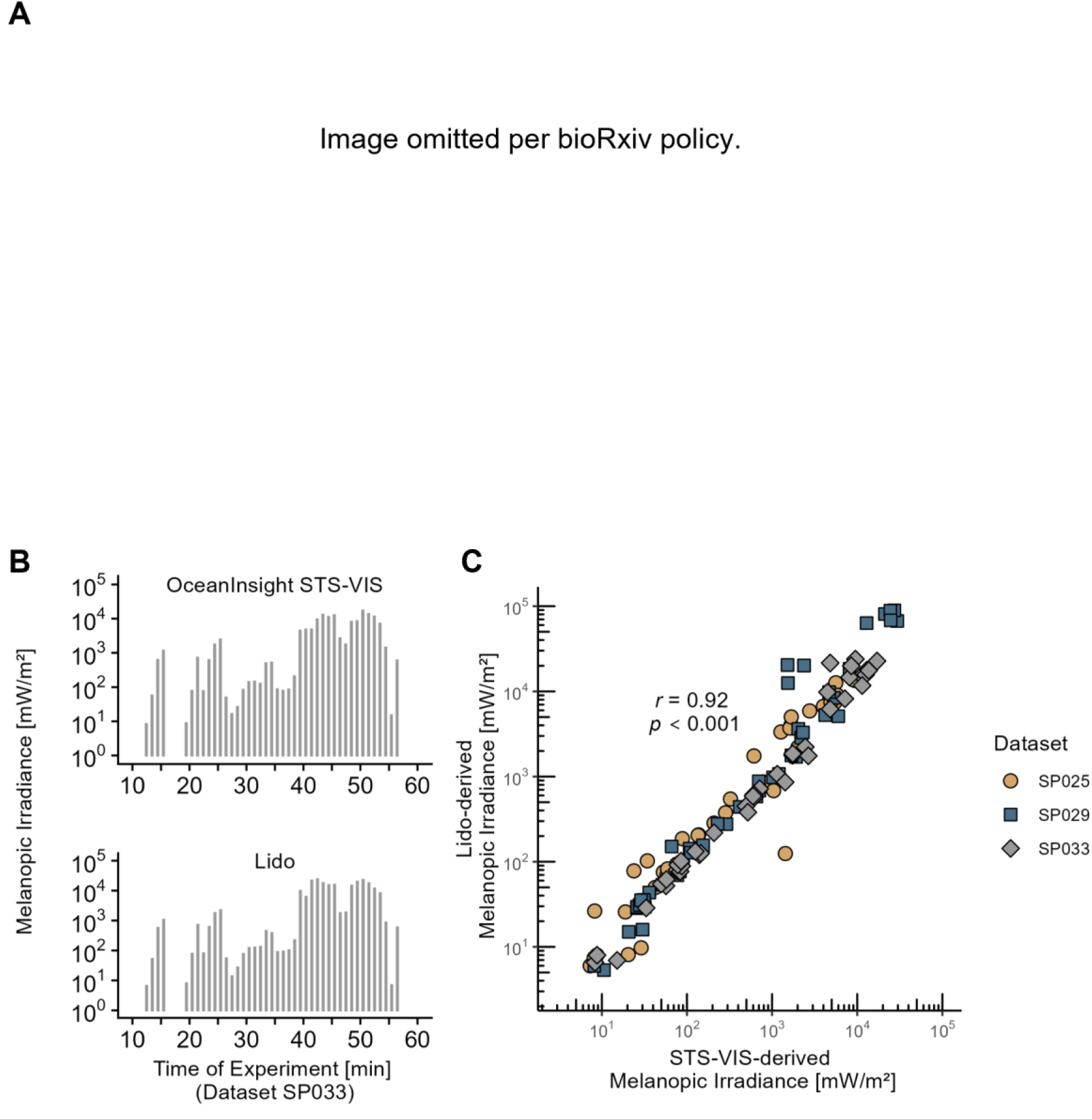
Cross-device comparison of light measurements. The Ocean Insight STS-VIS spectroradiometer used in the present data collection was compared to a light dosimetry device (lido) developed at Lucerne School of Engineering and Architecture ^74^. **A**, Colleague O.S. wearing the Ocean Insight STS-VIS spectrometer (forehead) together with the light-dosimeter (lido, left side of the head) in a room illuminated by blue LED light. **B**, Melanopic irradiance measured during the study protocol by the Ocean Insight STS-VIS (upper panel) and the comparison light dosimetry device (lower panel). **C**, Agreement of melanopic irradiance measurements (Pearson correlation) between the two devices, using three study protocol recordings (SP025, SP29 and SP033).

### Correction of timestamps

Due to an issue with the internal timekeeping of the Raspberry Pi microcontroller, the recorded timestamps (“Time” row in the uncalibrated spectroradiometric samples) and previously the file names were incorrect. To solve this problem, Python code was used to rename the recorded files according to their estimated start time (see the file “rename.py” in the “Python” folder of ^73^). Since each experimental trial started approximately ten minutes after the calibration video was recorded, the first sample of each trial was renamed to a timestamp ten minutes after the start of the calibration video (“Start date” and “Start time” from the “info.csv” files provided in the “gazecalibration/000” subdirectory). The following files were then renamed at 10-second intervals as they were recorded. In the current dataset, the filenames of both the calibrated and uncalibrated spectral irradiance samples and pupil images contains the corrected (estimated) timestamps of each sample.

#### Data quality checks

Like photographs, spectroradiometer measurements can become overexposed or “saturated” after rapid dark-to-light changes. In the present dataset, saturated spectrometer samples (410 files in total) were included as placeholders for completeness. “sat_files.csv” lists the corresponding CSV files containing all saturated/erroneous spectrometer samples. For the calibrated and interpolated spectra, the samples take the spectral irradiance value “0” at all wavelengths and should be considered missing and excluded when analysing the data.

In the processed pupil and light dataset ^70^ accompanying the Registered Report ^47^, participants with less than the threshold of 75% of data in the required processed data quality were excluded from the analysis, resulting in data from *N*=83 participants. Only these 83 participants are included in the present dataset ^48^, comprising 29,664 samples of paired pupil images and spectral irradiance in total. According to the condition phases of the experiment, the present dataset was categorised into “field data” (16,481 data pairs), “dark data” (6,013 data pairs), and “lab data” (5,994 data pairs). There were also residual transition samples (“mixed”, 1,176 data pairs) taken between the laboratory and field conditions and could not be clearly assigned to either condition. The metadata “Sample_phase_tab.pdf” of the current dataset ^48^ lists which samples (images and spectra) were taken in which phase of the experiment. The metadata file “sum_tab.pdf” lists the number of data pairs per phase and participant.

### Dark and thermal noise spectroradiometer calibration

Spectroradiometric measurements are affected by noise ^75^. To address this, some spectroradiometers subtract a matching dark spectrum generated by a shutter from each spectral sample collected. However, as the compact *Ocean Insight STS Vis spectrometer* did not feature a shutter mechanism, a calibration process was necessary using software after data recordings. The following dark and thermal noise calibration procedure has been described as part of a proceeding’s contribution ^76^.

For the *Ocean Insight STS Vis,* we found that the amount of noise in each spectral sample depended on the spectroradiometer board temperature at that time and the integration time (span) used to collect that sample. Specifically, a higher temperature and integration time led to elevated noise levels across wavelength pixels (see Fig. 5 **A**). During a sampling session, the board temperature generally increases, with the integration time for a sample being determined by the light level: Longer integration times are used in darkness, while shorter integration times are used in bright light. Our objective was to develop a dependable model to reliably predict “noise counts” [Z in number of sensor counts] from board temperature measured by an internal sensor [X in °C] and integration time [Y in ms]. We undertook this task corresponding to a previously developed calibration procedure ^75^.

**Figure 5.**
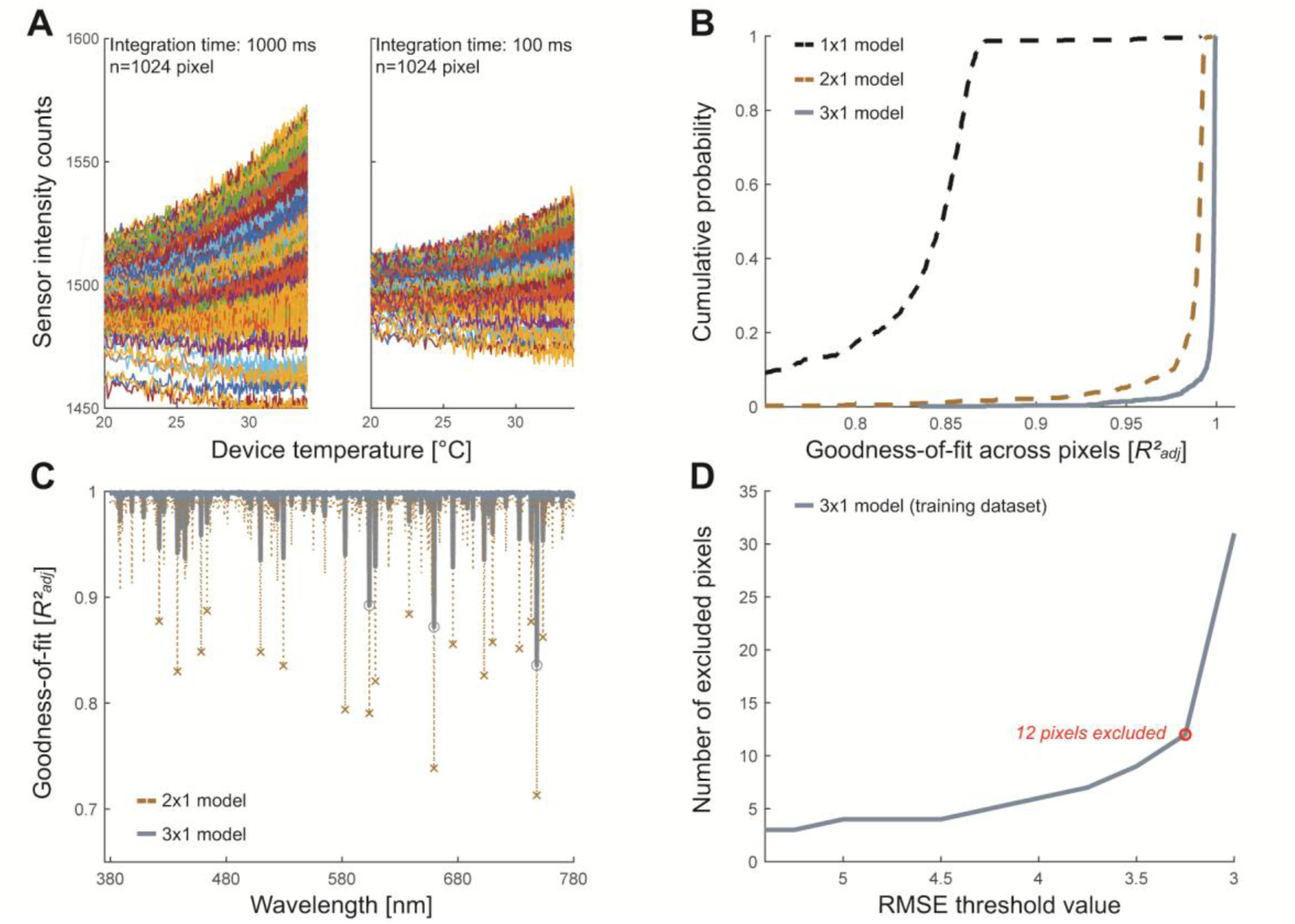
Dark and temperature noise calibration model selection. **A,** Variation of dark noise as a function of device temperature and integration time across all wavelengths. Each coloured line represents the sensor intensity count variation of a specific wavelength pixel measured by the spectroradiometer (STS-VIS, Ocean Insight Inc., Oxford, UK). The increase in both dark and thermal noise across wavelength pixels is more significant at higher device temperatures, particularly for samples taken with longer integration time (e.g. 1 sec compared to 0.1 sec). **B**, Calibration model comparison with cumulative probability of Goodness-of-fit. The plot illustrates the relative probability (y-axis) that a wavelength pixel in the respective polynomial model (indicated by the lines) takes a value equal to or less than the *R*²_adj_ value scaled on the x-axis. It displays the enhancement in goodness-of-fit from model 1×1 to 2×1 to 3×1. ***C***. Goodness-of-fit statistics per wavelength. Wavelength dependence of fit quality, contrasting the *R*²_adj_ values (y-axis) between model 2×1 (blue line) and 3×1 (dotted brown line) across wavelength pixels (x-axis). Pixels with *R*²_adj_<.9 are highlighted. **D**, Calibration pixel selection: visualisation of wavelength pixel exclusion in training dataset. *RMSE* denotes the Root Mean Square Error. Tightening the threshold beyond *RMSE*<3.25 leads to a sharp rise in the number of pixels to exclude (12 vs. 31 pixels).

#### Dark spectra database (training dataset)

A series of spectral measurements in darkness were collected with a plastic cap covering the light sensor of the spectroradiometer, incorporating different board temperatures and integration times in the range of our experimental conditions. The measurements were performed on a laptop running Windows 10 using the OceanView 2.0 software (Ocean Insight Inc., Oxford, UK) to operate the STS-VIS spectroradiometer. Six separate 60-minute sampling sessions were conducted, each with a specified integration time (10 ms, 50 ms, 100 ms, 500 ms, 1000 ms, 5000 ms). To manipulate board temperature, we stored the spectroradiometer in a refrigerator one hour before the session and allowed it to warm up during the session, resulting in a temperature range between ∼15 and ∼36°C, as measured by an internal sensor. In line with our experiment’s design, we sampled at 10-second intervals. During each of the 60-minute sessions, we collected 360 sensor count samples in darkness for every one of the 1,024 wavelength pixels measured by the spectroradiometer. Additionally, we collected 360 corresponding board temperature samples. Across all sessions, 2,160 raw count samples were obtained for each wavelength pixel, along with temperature measurements. This training dataset is part of the code repository ^73^ in the subdirectory “*Matlab\Calibration_files\baseline_data*”.

#### Polynomial curve fitting (training dataset)

Using the “fit” function from MATLAB’s Curve Fitting Toolbox™ (R2020a, The MathWorks, Inc.), we calculated goodness-of-fit statistics from five different polynomial models (1×1, 2×1, 2×2, 3×1,4×1*, see Table **2**) iterated over pixels in the visible wavelength range (847 pixels, 380–780 nm range) based on the training data set. Table **2** presents a summary of goodness-of-fit statistics for the five polynomial models on all pixels. The total variance of counts in a pixel remains unchanged across all models as we use the same raw data (training data), enabling comparison of models using their coefficient of determination statistics.

**Table 2.** Goodness-of-fit statistics for different polynomial models estimating dark and thermal noise. The numbers in the model notation (e.g. “3×1”) reflect the degrees of the independent variables used in each model. For instance, “3×1” indicates a model with independent variable X [temperature in °C] used in the third degree (cubic) and independent variable Y [integration time in ms] used in the first degree (linear). The coefficient of determination, *R²_adj,_* is adjusted for inflation by the number of coefficients in the model. Each pixel in the visible range of wavelengths (*N*=847) produced a corresponding *R²_adj_* value for its fitted polynomial model. Descriptive statistics for a single model were computed for all 847 pixels.

| Model degree | Number of Coefficients | Min $R^2_{adj}$ | Med $R^2_{adj}$ | Max $R^2_{adj}$ | n pixels, $R^2_{adj}<0.9$ |
| --- | --- | --- | --- | --- | --- |
| 1x1 | 3 | 0.108 | 0.848 | 0.991 | 837 |
| 2x1 | 5 | 0.713 | 0.990 | 0.997 | 18 |
| 2x2 | 6 | 0.716 | 0.990 | 0.997 | 18 |
| 3x1 | 7 | 0.836 | 0.999 | 0.999 | 3 |
| 4x1 | 9 | 0.844 | 0.999 | 1.000 | 3 |
| 3x2 | 9 | 0.840 | 0.999 | 0.999 | 3 |

#### Model selection (training dataset)

To determine the most suitable polynomial model, descriptive statistics of *R*²_adj_ values were utilised as the comparative metric. Typically, the coefficient of determination *R²* is defined as:

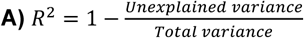

Unlike *R²* values, which increase by simply adding more predictors to a model, *R²_adj_* values adjust for this inflation effect. They achieve this by including a term that considers the number of predictors in each model relative to the total number of samples in the dataset ^77,78^.

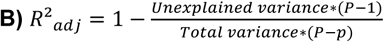

Where *P* denotes the number of samples in the training data set and *p* the number of coefficients in each respective model.

Table **2** and Figure 5 **B** and **C** demonstrate that meaningful improvements in model fit, as reflected in the *R²_adj_* values, were obtained by increasing the temperature from a first-degree (1×1) to a second-degree (2×1) to a third-degree (3×1) predictor. However, there were no additional improvements in goodness-of-fit by increasing integration time to a second-degree predictor (2×2) or by elevating temperature to a fourth-degree predictor (4×1). Consequently, the polynomial model “3×1” was chosen for calibration, resulting in the complete formula.

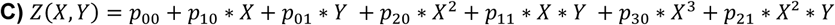

where *p* denotes the coefficients in the model, *X* is device temperature [°C], and *Y* is integration time [ms]

#### Pixel selection

To reduce misestimation in the noise calibration, we identified and excluded pixels that exhibited a high number of residuals in the prediction (“dead pixels”) in a two-step process. Given the high wavelength pixel resolution (847 pixels between 380 and 780 nm; ∼0.5 nm spacing) combined with the resampling to obtain evenly spaced 1 nm intervals, a reasonable level of pixel exclusion was achieved with minimal loss of information. In both stages, we utilised the Root Mean Square Error (RMSE) to indicate absolute deviation differences between the actual sensor count in the training dataset (Z) and the forecast produced by the 3×1 model (Ẑ). Opting for pixels based on their *R²_adj_* value wasn’t suitable since these values are calculated in proportion to the overall variance of sensor counts in each pixel of the raw data (see Formula B). Hence, *R*²_adj_ can yield high estimates simply due to large total variance while possibly also showing high residual values, which we want to prevent here. *RMSE* is given by:

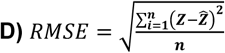

where *Z*_i_ is the i^th^ sample of sensor counts in darkness, *Ẑ* is the respective value predicted by the 3×1 model, and *n* denotes the number of samples taken for each pixel (in training dataset: *n*=2160; in validation dataset: *n*=67)

#### Pixel exclusion in training dataset

Our objective was to determine an appropriate *RMSE* threshold for the training dataset while keeping pixel exclusion-related data loss in check. We plotted different *RMSE* threshold levels, starting from *RMSE* ≥ 10 [sensor counts], against the resulting number of pixels to exclude in a stepwise decrease of 0.25. Figure 5 **D** illustrates a sharp increase in the number of pixels to be excluded when the threshold decreases from *RMSE* ≥ 3.25 (12 pixels) to *RMSE* ≥ 3.0 (31 pixels), where the line indicating the “*number of pixels to exclude*” in the training data forms an ‘elbow’. As a result, the 12 pixels with *RMSE* ≥ 3.25 in the training dataset were removed from future analysis.

#### Pixel exclusion in the validation dataset

The remaining 835 pixels were subsequently employed to predict noise counts from an additional 67 dark samples collected during an early test run of the experiments. This “validation dataset” is available as part of the code repository ^73^ in subdirectory “*Matlab/Calibration_file/pixelselection_dataset*”. The *RMSE* threshold was not set as rigidly as before since these dark samples were obtained in an unlit room, unlike the ones which were previously covered with a plastic cap and yielded the maximum 5-second integration time, consequently resulting in a higher noise rate. Upon inspection by eye, conspicuous pixels were once again identified following the computation of corresponding *RMSE* values. Three additional pixels generating an RMSE value greater than 25 were identified and subsequently excluded from further analysis.

#### Noise spectra computation

In the final 832 wavelength pixels, the seven coefficients (*p00* to *p21*) of the (3×1) model according to formula C were used to predict a “noise count” from the board temperature [X] and integration time [Y] samples corresponding to each light sample in the experiment. The calibration process was concluded by subtracting the “noise spectra” from the empirical spectral samples during early data pre-processing.

All MATLAB code used during the dark and thermal noise spectroradiometer calibration described above is available in the “Matlab” folder of the code repository on GitHub ^73^. This includes code to generate the various models, tables and figures of the dark and thermal noise calibration file, as well as the training and validation datasets, on which this analysis was based.

## Usage notes

### Processing steps in the prior report

In our prior Registered Report publication ^47^ using processed data ^70^ (based on the current raw dataset ^48^), we set out to test how age and different light levels affect pupil size under naturalistic, real-world conditions and whether melanopic quantification of light predicted steady-state pupil size better than photopic illuminance under these conditions. We further explored whether there is evidence for sex, iris colour or reported caffeine consumption affecting steady-state pupil size under real-world conditions. An example of the age effect on pupil size is given in Figure **6**, which shows typical sets of processed pupil size data from a young (18 years old, left panel), middle-aged (44 years old, middle panel) and old (80 years old, right panel) participant plotted as a function of the log_10_-transformed melanopic equivalent daylight illuminance (mEDI) derived from spectral irradiance data.

**Figure 6.**
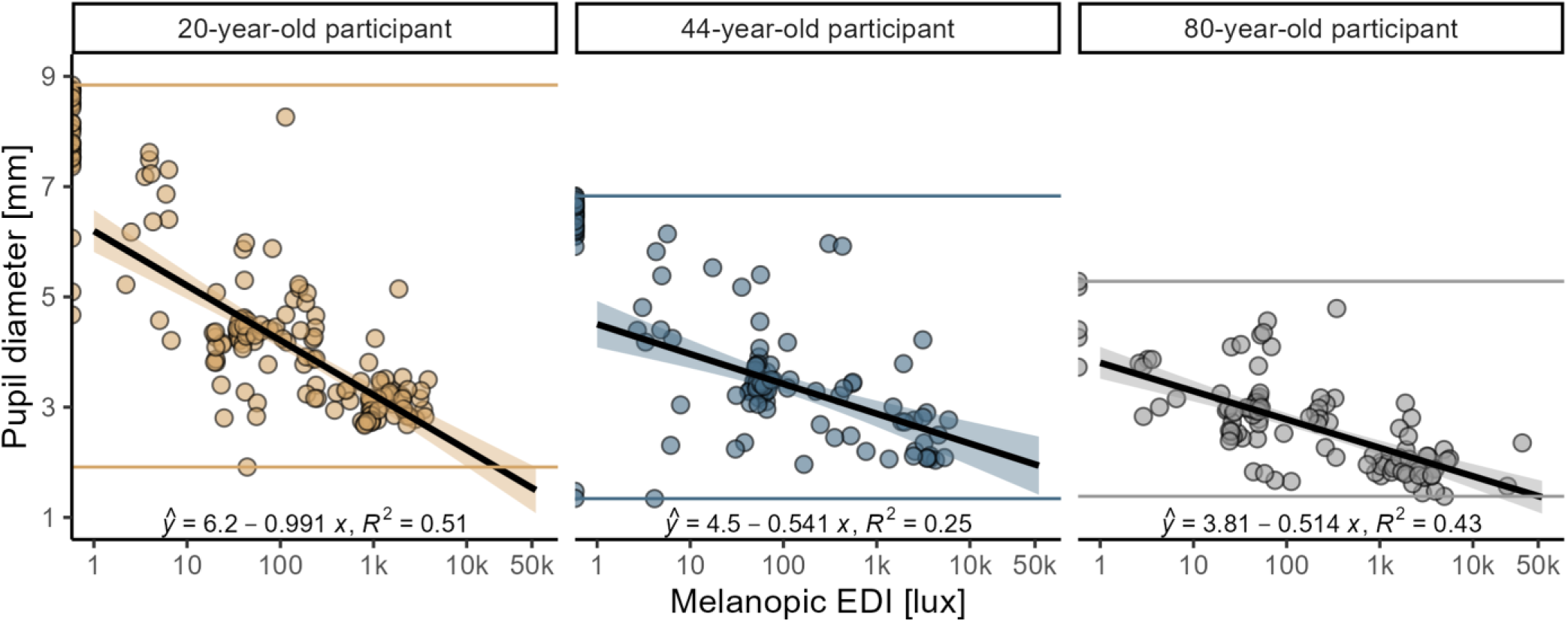
Age comparison of three typical sets of processed pupil size data. Pupil diameter (mm) is given as a function of melanopic equivalent daylight illuminance (mEDI; lux) derived from the spectral irradiance data. The scatterplots show data from the field and dark adaptation conditions of the experiment in three different subjects: a young (20 years old, left panel), a middle-aged (44 years old, middle panel) and an old (80 years old, right panel) participant. The regression line and equation show the linear relationship between log_10_-scaled mEDI values in lux and pupil diameter in mm. The coloured horizontal lines mark the minimum and maximum pupil diameter values for each subject.

A detailed description of the workflow and R Code processing steps used to conduct data analysis in the Registered Report ^47^ is provided in that paper and the readme file of the corresponding GitHub repository ^58^. A brief description of the data processing is provided in the following.

#### Pupil size estimation

For the processed data of the Registered Report ^47^, the still eye images (Fig. **1B**) were pasted together as a sequence per participant and subjected to the intrinsic three-dimensional model derived in the calibration phase by means of Pupil Labs software (release version 1.15.71, https://github.com/pupil-labs/pupil/releases), resulting in pupil diameter data in millimetres along with a pupil detection confidence. The settings used during the pupil extraction process in the Pupil Capture software ^71^ of each participant was recorded in a screenshot (*<Participant ID>_PLsettings.png., see Data Records)* and provided in each participants’ subfolder (e.g. “*SP001*”) of the present dataset ^48^. The screenshot shows a visualisation of the 3D pupil model ^45^ including the *pupil intensity range*, which gives “*the minimum darkness of a pixel to be considered as the pupil*” ^72^. This pupil intensity range value took the value 10 at default and was adjusted (minimized) for every participant’s data so that the pupil was always fully covered while having as little leakage outside of the pupil as possible. Differences between participants occurred mainly due to differences in iris colour and differences in illumination of the eye during the field condition. The screenshot also contains “*Pupil min*” (10) and “*Pupil max*” (100) and “*Model sensitivity*” (0.9970) values, which were kept consistent across participants. Low pupil detection confidence values (<0.6, according to Pupil Labs guidelines) occurred when the pupil size estimation was complicated or disrupted due to extreme viewing angles or partial occlusion by eyelashes in the eye images, as participants were performing real-world tasks and, hence, blinking, squinting, and moving their eyes.

#### Spectral data processing

From the calibrated and interpolated spectral data, α-opic irradiance values were calculated using spectral sensitivity curves from the CIE S026/E:2018 standard ^18^, alongside photopic illuminance in lux based on the CIE 1924 photopic luminous efficiency curve ^79^ and additional colour characteristics such as the correlated colour temperature (CCT (K) - Ohno, 2013 ^80^), Colour Rendering Index (CRI) ^81^, and CIE 1931 xy chromaticity coordinates (2° observer) ^82^. These light characteristics can, for example, be generated by using the luox application ^25^ or the CIE toolbox ^23,24^ on the calibrated and interpolated spectra of this dataset.

#### Data quality checks in processed data

To ensure high-quality data for the pre-registered hypotheses tests, the following data quality checks were applied to the processed data for the Registered Report ^47^: 3D model pupil size estimates that exceeded the defined maximum pupil range (<1 or >9 mm; 0.16% in total) were set as missing. 3D model pupil size estimates with low detection confidence (<0.6 according to Pupil labs software), lacking a good quality 3D model fit, for example, due to due to extreme angles of gaze, partial covering by eye lashes or light reflections on the pupil, were set as missing values. This comprised 43.37% in total in the processed data. Finally, light data derived from saturated/ erroneous spectroradiometer samples were set as missing (1.41% of samples).

Giving contextual information on the processed light and pupil data, Figure **7** shows the autocorrelation of processed light data from the Ocean Insight spectroradiometer and the estimated pupil size data with a 3-minute lag. The high autocorrelations show relatively constant real-world light conditions, mirroring the data collection protocol, where participants stayed in each light situation for at least 60 seconds.

**Figure 7.**
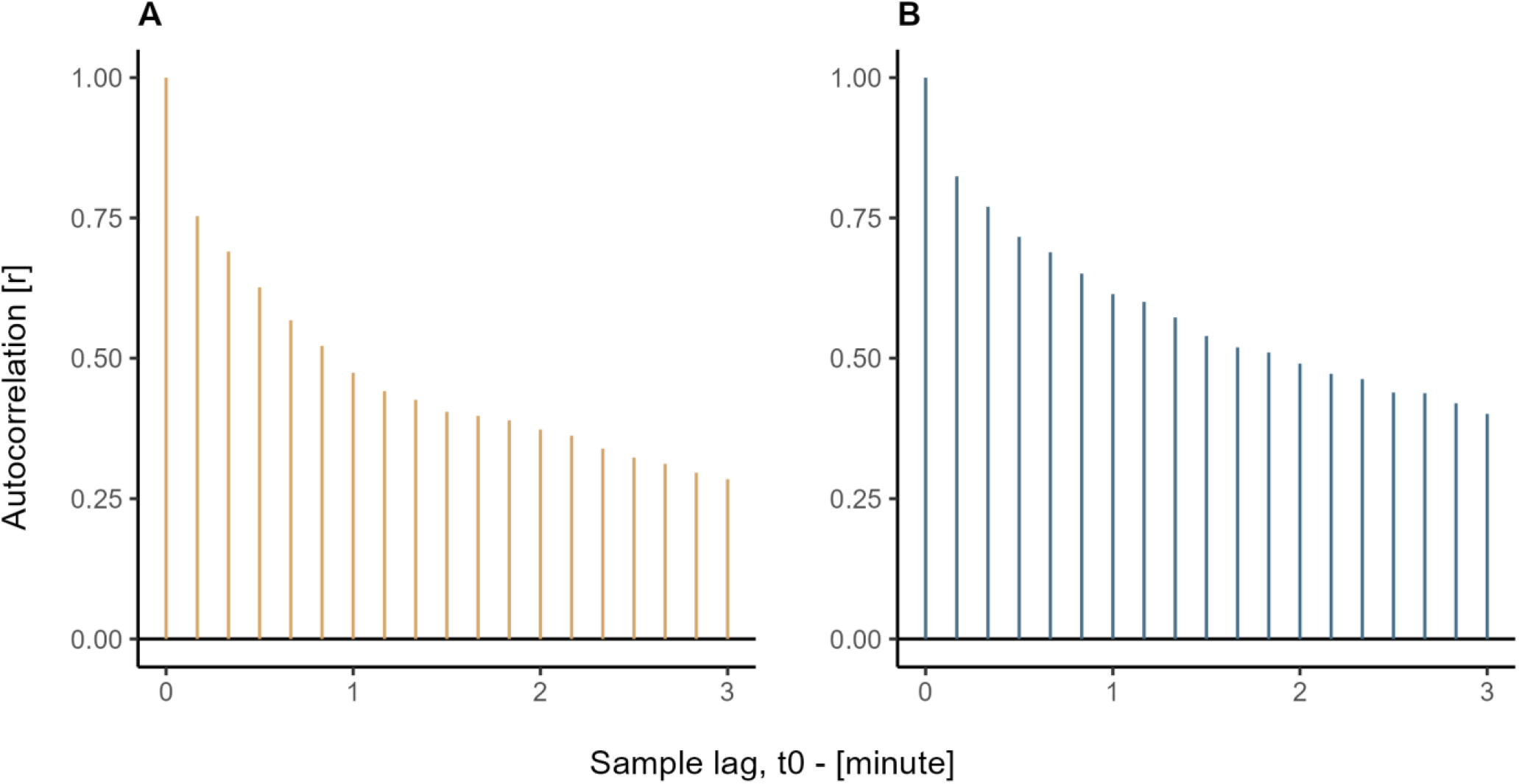
Autocorrelation of processed light and pupil data from the field condition. **A**, Autocorrelation of melanopic equivalent daylight illuminance (mEDI) derived from the spectral irradiance measurements across a 3-minute time lag during the field conditions of the protocol (*N*=83). **B**, Autocorrelation of pupil diameter derived from the pupil images across a 3-minute time lag during the field conditions of the protocol (*N*=83). Concurrent samples (*t0*) of pupil size and mEDI were highly autocorrelated with the sample 10 seconds prior (*r*>0.75, Pearson coefficient) and remained slightly autocorrelated (*r*>0.25, Pearson coefficient) with the sample 3 minutes prior.

### A foundational data set for future scientific work

Beyond prior analyses of the data set, we see the following future use cases for this dataset:

#### Development of novel image-based methods for pupil detection

The dataset contains a large number of infrared images of the human pupil, recorded under varied natural lighting conditions and from a diverse group of participants across the lifespan. This resource can support the testing and improvement of algorithms for detecting and measuring pupil size with greater accuracy. Reliable pupil detection methods are essential for advancing research in vision science and clinical applications.

#### Characterization of statistical regularities in real-world scenes

The spectral data in this dataset reflect the diversity of real-world lighting conditions. These data can be analysed to better understand patterns in natural and electric light exposure. This information can be used to study the relationship between light environments and human vision and may be relevant for applications in lighting design and exposure assessment.

#### Development of novel spectrally resolved methods for predicting pupil size

The dataset combines pupil size measurements with spectral irradiance data across different age groups, providing an opportunity to study how light spectra influence pupil size. This can support the development of methods to estimate pupil size based on spectral data, considering factors such as age and individual variability. These insights are useful for studying retinal light exposure and its consequences.

## Code availability

The software code to drive the acquisition device used to generate the presented raw dataset is proprietary and not available as part of this dataset. However, Python code used to correct timestamp-based filenames, MATLAB code to generate the dark and temperature noise calibration, and R code to generate metadata files, figures and tables of the present data descriptor are publicly accessible under the MIT licence in a GitHub repository ^73^ published together with this data descriptor. The current raw dataset, together with metadata, is available on figshare ^48^.

For the processed dataset analysed in the Registered Report ^47^, all code and data are publicly accessible. Code and processed data are available on GitHub ^58^ under the MIT license. Processed data and the laboratory logbook are available under the CC-BY 4.0 license on figshare ^70^. Other supporting materials are available on figshare ^49^ under the CC-BY 4.0 license.

## Ethical approval

Ethical approval has been granted from the cantonal ethics commission (Ethikkommission Nordwest- und Zentralschweiz, project ID 2019-01832, see Supporting materials on figshare ^49^). Mandatory measures to prevent the spread of COVID-19 in Switzerland were implemented during data collection.

## Acknowledgements

We thank Josefine Degen, Ann-Sophie Fiechter, Cielle Hawrylenko, Majlinda Maliqi, Aurora Monticelli, and Sebastian Saraceno for their assistance in the data collection, Dr Brian Cardini for his support in writing the demographic analysis code and Elias Mattern for his support in writing the python code for correcting the timestamps of the raw data.

During parts of this work, M.S. was supported by a Sir Henry Wellcome Trust Fellowship (Wellcome Trust 204686/Z/16/Z) and a Junior Research Fellowship from Linacre College, University of Oxford. R.L. was funded by the European Training Network LIGHTCAP (project number 860613) under the Marie Skłodowska-Curie actions framework H2020-MSCA-ITN-2019 and by the Nikolaus and Bertha Burckhardt-Bürgin Foundation. The data collection of this study was funded by an investigator-initiated research project supported by Ocean Insight.

## Author contributions

The authors made the following contributions.

Rafael Robert Lazar: Conceptualization, Data curation, Formal Analysis, Investigation, Methodology, Project administration, Supervision, Visualization, Validation, Writing – original draft, Writing – review & editing;

Manuel Spitschan: Conceptualization, Data curation, Funding acquisition, Methodology, Project administration, Resources, Supervision, Visualization, Validation, Writing – original draft, Writing – review & editing.

